# Obtaining polygenic transcriptome risk scores (PTRS) directly from GWAS summary statistics

**DOI:** 10.1101/2022.07.10.499509

**Authors:** Yanyu Liang

## Abstract

Polygenic Transcriptome Risk Scores (PTRS) are variations of Polygenic Risk Scores (PRS) that use genetically predicted transcriptome as features for prediction instead of directly using genetic variants. We have shown that when PTRS is combined with PRS, they can yield improved prediction performance and portability across populations (Liang et al., 2022). Given the difficulty of training PTRS using large scale individual-level data (due to both computational burden and the lack of data access), we developed a user friendly software that infers PTRS using GWAS summary results and reference LD. We tested three summary statistics-based PTRS approaches: i) Clumping and thresholding (clump-PTRS), keeping trait associated genes while removing highly correlated ones; ii) Summary statistics-based elastic net PTRS (S-EN-PTRS), an extension of lassosum (Mak et al., 2017) to predicted transcriptome; iii) Naive-PTRS, the sum of predicted expressions of significantly associated genes weighted by PrediXcan-estimated effect sizes (Gamazon et al., 2015). Despite reports that individual-level trained elastic net PTRS outperformed clump-PTRS in (Liang et al., 2022), for most of the 11 traits used in the comparison, clump-PTRS outperformed S-EN-PTRS, which outperformed naive-PTRS.

## Introduction

PRS is a promising tool to translate GWAS discoveries into clinically-relevant preventive and therapeutic strategies, but there are some challenges that need to be addressed. One issue with PRS is the loss of performance when applied to populations that are underrepresented in current GWASs, i.e. non European-descent groups (Martin et al., 2019). To address this problem, we have proposed PTRS, which uses genetically predicted trancript levels instead of genetic variants as predictors of complex traits (Figure 1B) (Liang et al., 2022). We have shown that PTRS can help improve portability across populations and when combined with traditional PRS, they can yield better performance than PRS alone (Liang et al., 2022).

**Figure 1:**
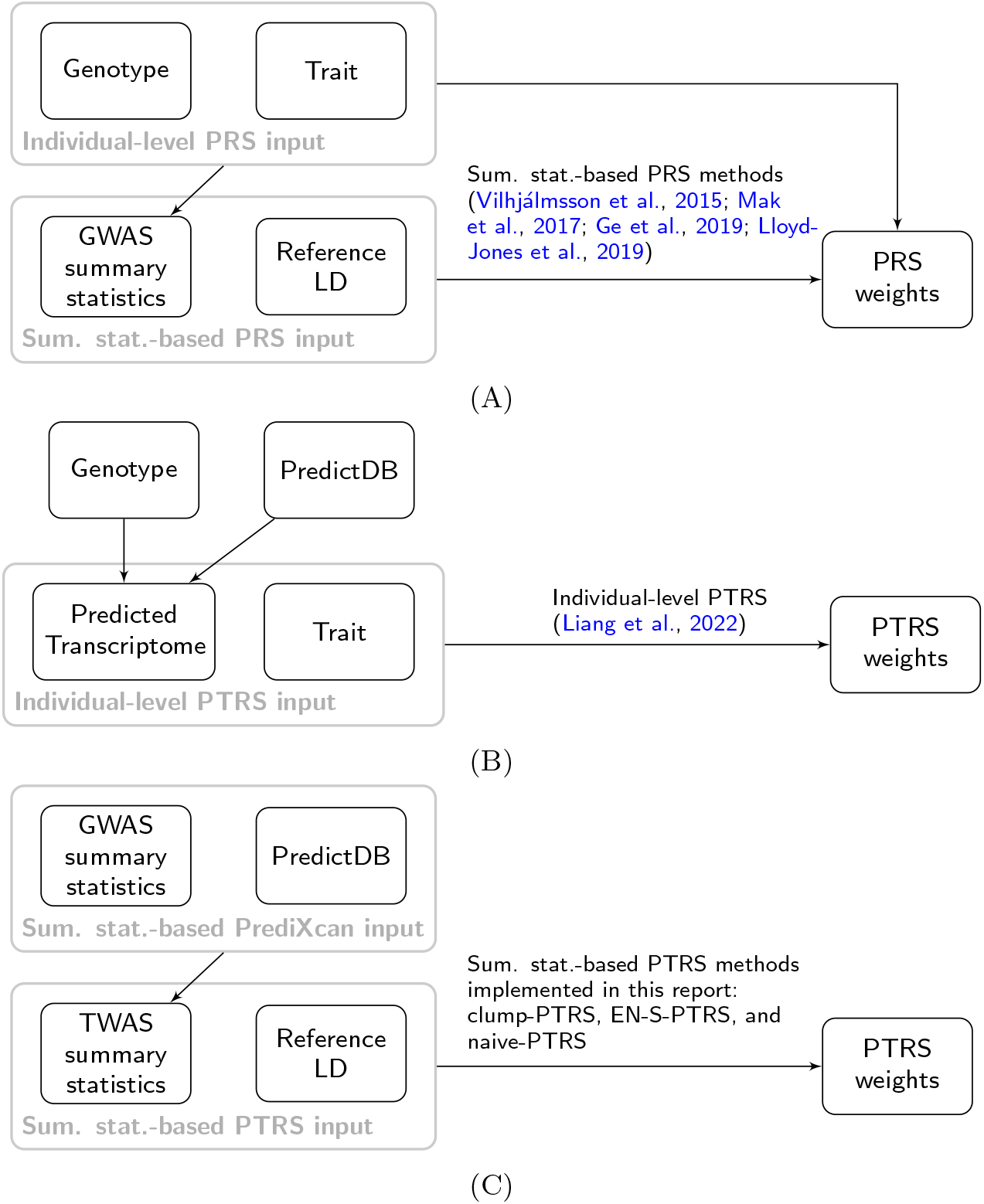
Overview of PRS and PTRS workflows. **(A)** PRS weights are calculated typically from the GWAS summary statistics and a reference LD (correlation between genetic variants) by summary statistics-based PRS methods (Vilhjálmsson et al., 2015; Mak et al., 2017; Ge et al., 2019; Lloyd-Jones et al., 2019); **(B)** PTRS weights are calculated by (Liang et al., 2022) using individual-level genotype and trait data in the UK Biobank cohort. It first calculates the predicted transcriptome of each individuals from genotype and prediction models of gene expressions (i.e. PredictDB from predictdb.org) and then obtains PTRS weights from predicted transcriptome and trait data with a mini-batch elastic net solver tailored to biobank-scale dataset; **(C)** The summary statistics-based PTRS approach implemented in this report uses TWAS summary statistics (obtained from GWAS summary statistics, PredictDB, and a reference LD by summary statistics-based PrediXcan/TWAS (Gusev et al., 2016; Barbeira et al., 2018)) and a reference LD to derive PTRS weights.

Another issue with PRS is that very large training sets (GWASs with very large sample sizes) are needed to obtain relevant prediction accuracy. However, obtaining access to and handling large-scale genetic and phenotypic data can be challenging due to computational and regulatory hurdles. Fortunately, PRS weights can be derived directly from the summary results of GWAS in combination with reference LD information (Figure 1A) (Vilhjálmsson et al., 2015; Mak et al., 2017; Ge et al., 2019; Lloyd-Jones et al., 2019).

In this report, similar to the essence of summary statistics-based PRS approaches, we develop a tool that derives PTRS using GWAS summary statistics and a reference LD (Figure 1C). We implement several approaches to derive PTRS and provide a user-friendly software https://github.com/liangyy/SPrediXcan2PTRS.

## Results

To calculate PTRS based on GWAS summary statistics, we extended three common approaches used for PRS construction using GWAS summary statistics: i) clumping and thresholding, ii) summary-based elastic net as implemented in lassosum (Mak et al., 2017), and iii) simple significance thresholding. Clumping and thresholding consists of selecting the most significant features and filtering out features that are highly correlated with each other or do not pass some significance threshold. The lassosum infers the elastic net trained predictors using GWAS summary statistics (Mak et al., 2017). Simple significance thresholding consists of filtering out the variants with non-significant GWAS p-values and using the GWAS effect sizes as the weights in the PRS. We refer to these extensions as clump-PTRS, S-EN-PTRS, and naive-PTRS, respectively.

All three summary statistics-based PTRS approaches start by calculating the TWAS summary statistics using the GWAS summary statistics and transcriptome prediction weights (PredictDB in Figure 1). Clump-PTRS removes genes that are highly correlated and those that do not pass a given p-value threshold. S-EN-PTRS is an implementation of lassosum with TWAS summary statistics taking the place of the GWAS summary statistics. Both the clump-PTRS and S-EN-PTRS require information on the correlation between SNPs which are involved in the prediction of gene expression levels. Naive-PTRS uses the z-scores (or effect sizes) from the TWAS summary statistics as weights for the PTRS. Further details are described in Methods.

To examine the performance of the summary statistics-based PTRS approaches, we selected 11 quantitative trait GWASs listed in Table 1 and calculated the PTRS weights using each of the three approaches described above (Methods). We evaluated the performance of these PTRSs using 5,000 randomly sampled European-descent participants in the UK Biobank. We calculated the Spearman correlation between PTRS scores and the observed UK Biobank traits (Table1) as a measure of the performance of the PTRS (Methods). For the S-EN-PTRS we used a range of shrinkage values (0.01, 0.1, 0.3, 0.5, 0.7, 0.9) and set the mixing parameter *α* = 1 (effectively a lasso penalty was used). For the clump-PTRS, we used a squared correlation cutoff of 0.1 and a sequence of p-value cutoffs (applied after clumping): 10^−7^, 10^−6^, 10^−5^, 10^−4^: 10^−3^, 0.005, 0.01, 0.05, 0.1, 0.5, 1.

**Table 1:**
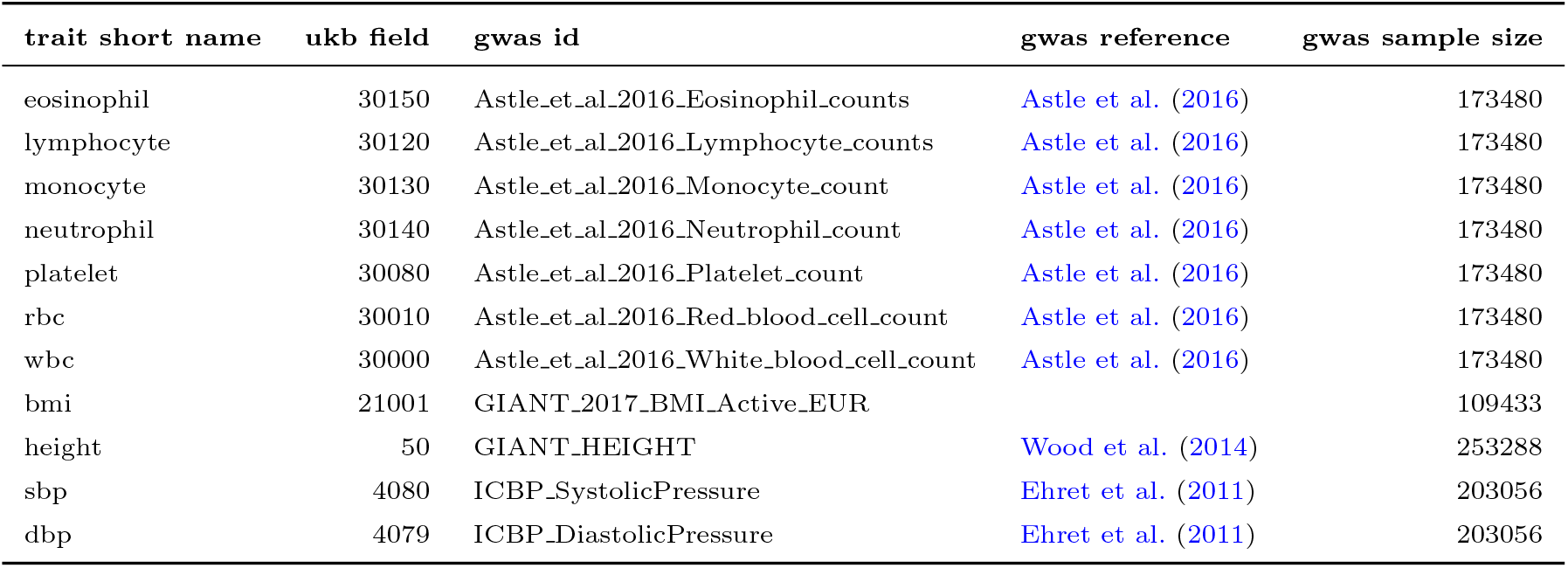
Information on the 11 quantitative traits being used for examining the performance of PTRS. We used the GWAS harmonized in Barbeira et al. (2021). Column “gwas id” indicates the GWAS ID used in Barbeira et al. (2021). Column “gwas reference” shows the reference to the GWAS. Column “ukb field” shows the Field ID in the UK Biobank

As shown in Figure 2, among most of these 11 quantitative traits, clump-PTRS performed the best, followed by S-EN-PTRS with the performance increasing with higher shrinkage parameters. Naive-PTRS showed the lowest performance.

**Figure 2:**
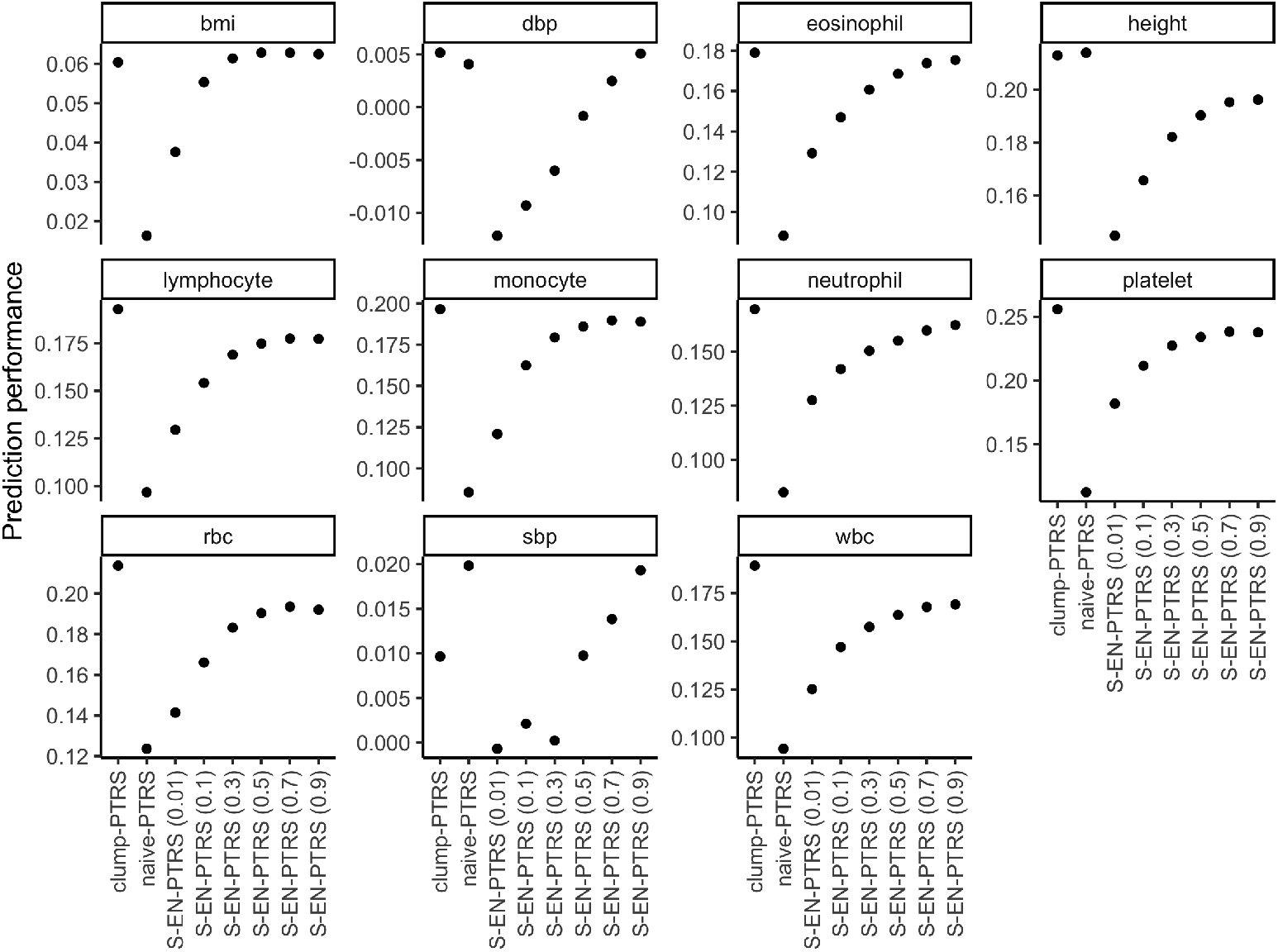
Performance of S-EN-PTRS, clump-PTRS, and naive-PTRS. Different summary statistics-based PTRS approaches are shown on x-axis. For S-EN-PTRSs, the value in the parentheses indicates the offset term *o* being used. PTRSs were evaluated using 5,000 individuals in UK Biobank where half of the samples were used to determine the best-performing hyperparameters (for clump-PTRS, it is the p-value threshold; for S-EN-PTRS, it is the mixing parameter λ). The y-axis shows the Spearman correlation on the other half of the samples with demographic covariates and genetic PCs being adjusted (more details in Methods).

The lower performance of S-EN-PTRS was unexpected given that the individual-level elastic net-based PTRS outperformed clump-PTRS in our previous study (Liang et al., 2022). One possible explanation of this low performance is the difference in LD between the source GWAS study and the reference one used here. Moreover, we observed that S-EN-PTRS performance increases as the offset term *o* becomes larger (see definition of the offset in Methods). Since the offset term measures the tendency to treat all genes as independent predictors instead of taking LD information into account, this result also suggested that S-EN-PTRS is potentially sensitive to LD panel.

We provide an out-of-box implementation of the three summary statistics-based PTRS approaches, clump-PTRS, S-EN-PTRS, and naive-PTRS, under a unified interface. It takes S-PrediXcan results along with a reference LD panel and generates the corresponding PTRS weights. The software is available at https://github.com/liangyy/SPrediXcan2PTRS.

## Discussion

In this work, we implemented three summary statistics-based PTRS approaches. We briefly examined the performance of these summary statistics-based PTRSs on 11 quantitative traits while using out-of-sample LD panel. We observed that, with out-of-sample LD panel, clump-PTRS outperforms S-EN-PTRS even though it’s been reported that EN-PTRS outperforms clump-PTRS when using individual-level data (Liang et al., 2022). Such discrepancy in performance might be attributed to the LD mismatch since, comparing to clump-PTRS, S-EN-PTRS is more sensitive to the accuracy of the LD panel. To support this claim will require a comprehensive survey of the behavior of these approaches under different scenarios (e.g. variations on the qualities of LD panels, GWASs, and gene expression prediction models). However, a deep performance survey is beyond the scope of this report and we defer it to future work.

Nonetheless, on the basis of the brief performance comparison results in this report, we caution that one should use S-EN-PTRS carefully with an out-of-sample LD panel. With mismatched LD, one could increase the offset parameter in S-EN-PTRS to reduce the effect of mismatch LD on the prediction performance. In practice, one could include both S-EN-PTRS and clump-PTRS as part of their model selection and hyperparameter tuning scheme.

We hope this report along with the software could ease and accelerate the future study and development of PTRS.

## Methods

### Details of S-EN-PTRS implementation

For *N* samples and *K* genes, let *y* ∈ ℝ^*N*×1^ represent the phenotype and let *G* ∈ ℝ^*N*×*K*^ be the predicted expression matrix. As described in (Liang et al., 2022), EN-PTRS is defined by the following problem:

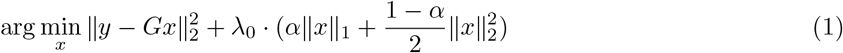

The EN-PTRS weights correspond to the solution to Eq 1.

We note that the amount of the regularization on each gene depends on the scaling of the corresponding column in *G*. To resolve such ambiguity, we further assume that each column of *G* is standardized and so is *y*. With this assumption, implicitly, the gene-level effect of gene *k* (*x*_*k*_) quantifies the amount of phenotypic increase (in the unit of standard deviation of the phenotype) under one unit increase of the predicted expression level of gene *k* (e.g. the predicted expression goes from the population mean to the population mean plus standard deviation).

Under this scaling, Eq 1 is equivalent to

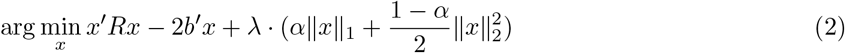

*R* is the sample correlation matrix of *G* and 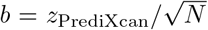. Since, equivalently, we use GWAS cohort for EN-PTRS training, *N* is essentially the GWAS sample size. And *λ* = *λ*_0_*/N*.

In practice, we don’t have access to *R* (the in-sample sample correlation). Instead we calculate the sample correlation using a reference LD panel as 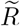. When the GWAS cohort and the reference LD panel are from the same ancestry group, 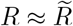 especially when both GWAS and reference LD panel have a lot of samples. In practice, to stabalize the model fitting (similar to lassosum (Mak et al., 2017)), we add an offset term to 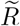 when approximating *R*. In other words, we let 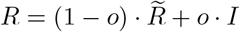.

We treat *o* ∈ (0, 1), *α* ∈ [0, 1], and *λ* ∈ (0, ∞) as hyperparameters. For a given pair of *o* and *α*, we determine the maximum *λ* by solving for the minimum possible *λ* satisfying the KKT condition when *x* = 0 is the solution ((Friedman et al., 2010)). In other words,

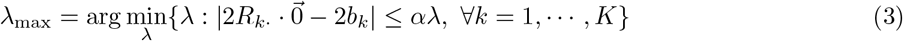

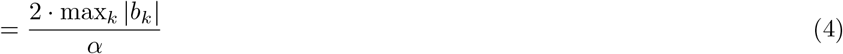

We solve for a sequence of equally spaced (in log-scale) *λ* values from *λ*_max_ to *λ*_min_ and usually *λ*_min_ = *λ*_max_*/*100.

Similar to (Friedman et al., 2010), we solve Eq 2 by coordinate descent. For the sequence *λ*_1_ = *λ*_max_, *λ*_2_, ⋯, *λ*_*J*_ = *λ*_min_, we first set 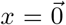 for *λ* = *λ*_max_. And then, for each *λ* = *λ*_*j*_, we initialize *x* as the solution from *λ* = *λ*_*j*−1_.

Additionally, to obtain 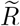, we need information more than the one required for S-PrediXcan, since we need to know both the variant/variant covariance for each variant pair within a gene and the variant pair from two different genes. We note that it is not suitable to pre-compute gene covariance since, usually, the GWAS may miss some genetic variants in the prediction models and we should exclude these variants for the PTRS training. Instead, similar to the S-PrediXcan implementation, we calculate 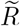 on the fly using the pre-computed variant/variant covariance from the reference LD panel. In practice, we consider gene/gene covariance for each chromosome and assume the covariance between genes from different chromosomes equals to zero. With this assumption, we need to pre-compute and store variant/variant covariance for all the variants that appear in at least one of the prediction models and lie in the same chromosome. For chromosomes 1 to 22, the number of variants to consider ranges from 4,000 to 30,000, for which we need to store all the pairwise information. In fact, the number of samples in the reference LD panel *N*_LD_ is usually smaller than the number of variants being considered *N*_SNP_ and it is an order of magnitude smaller when using GTEx data as the reference (sample size is about hundreds) The *N*_SNP_-by-*N*_SNP_ variant/variance covariance matrix can be represented by only *N*_LD_ eigenvalues and eigenvectors. We make use of this property and save the eigenvalue decomposition of the variant/variant covariance matrix instead.

In principle, we need to loop over the genome-wide variants until convergence. But here, we also implement an alternative scheme in which we loop over variants within each chromosome (this is similar to lassosum (Mak et al., 2017) which solves elastic net models for each LD block while using the same *y* for all blocks). We note that this chromosome-wise approach usually gives similar results as the genome-wide iteration and converges a little faster.

### Details of clump-PTRS implementation

Given a list of genes with PrediXcan z-scores and *r*^2^, a cutoff on squared correlation between genes, we want to perform clumping such that genes with duplicated signals (genes which are “highly” correlated with a more significant gene) are removed. The procedure is similar to (Liang et al., 2022) and it is outlined in Algorithm 1.

#### Algorithm 1

Clumping procedure

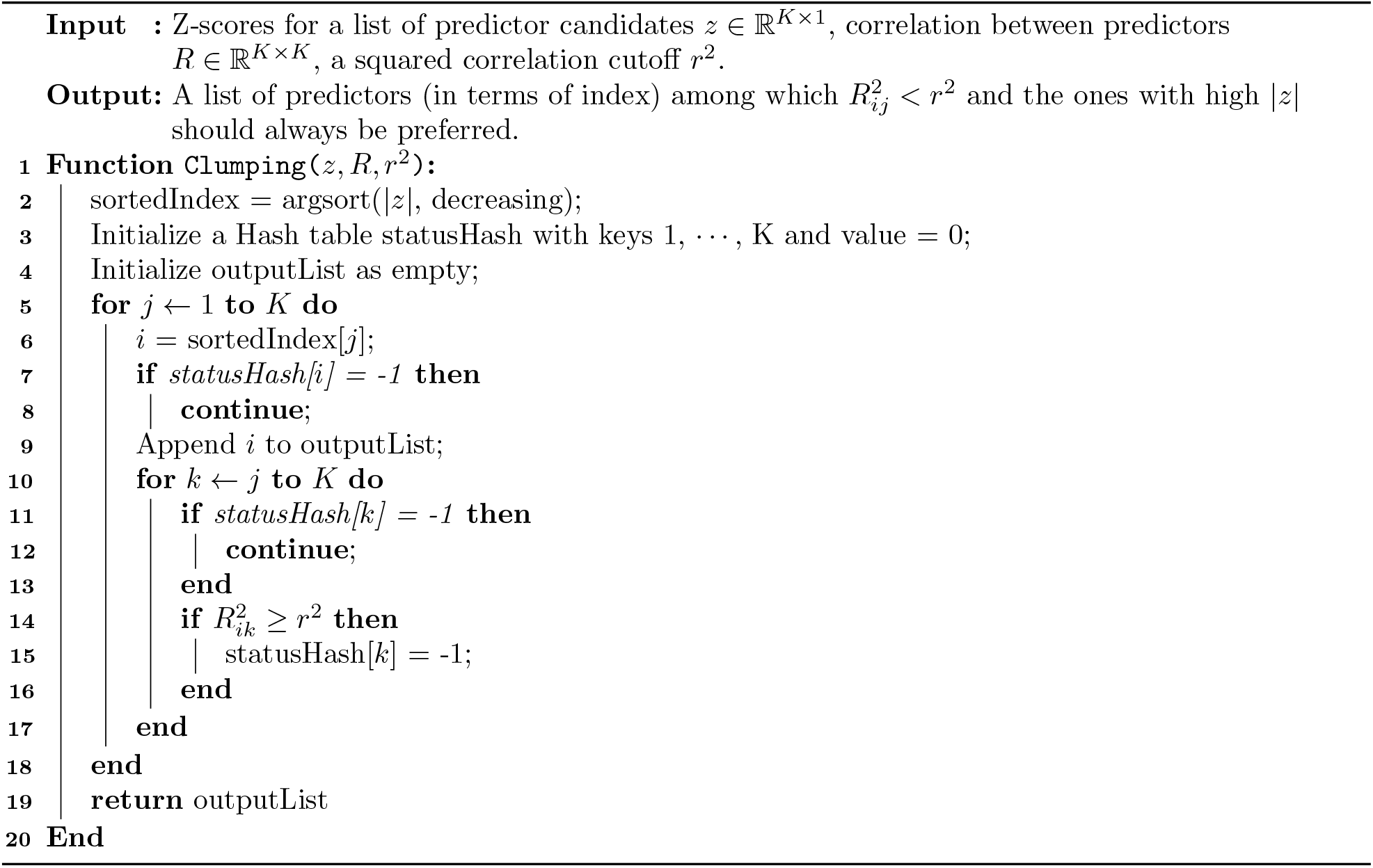

### Applying summary statitics-based PTRS to 11 GWASs

We applied clump-PTRS, S-EN-PTRS, and naive-PTRS to 11 GWASs listed in Table 1. These GWASs were harmonized previously and imputed in GTEx v8 WGS samples (Barbeira et al., 2021). We used the CTIMP models and cross-tissue gene expression imputation models (Hu et al., 2019) from Whole Blood, which were trained on GTEx v8 European samples (Barbeira et al., 2020). The variant/variant covariance was obtained from GTEx v8 European samples.

### Evaluating PTRS performance using UK Biobank data

We evaluated the prediction performance by calculating the Spearman correlation between the predicted and observed phenotype values on 5,000 randomly selected UK Biobank participants with European ancestry. To take covariates into account, we regressed out the top 10 genetic PCs, age, and sex (and second order terms of age and sex) from both the observed and the predicted values before calculating the correlation. The corresponding UK Biobank phenotypes used for the evaluation are shown in Table 1.

To select the hyperparameters, we split the selected participants into two halves. We used the first half samples to determine the hyperparameters (*λ* for S-EN-PTRS and the p-value cutoff for clump-PTRS) by selecting the best-performing models. And then we used the second half of the samples to calculate the prediction performance under the models with the selected hyperparameters.

## Code Availability

The software implemented for this work is available at https://github.com/liangyy/SPrediXcan2PTRS.

## Acknowledgment

We thank Hae Kyung Im for helpful discussion. We thank Natasha Santhanam and Beau Burnett for helping us edit the report. This research has been conducted using the UK Biobank Resource under Application Number 19526.

